# CD24 regulates the formation of ectosomes in B lymphocytes

**DOI:** 10.1101/2024.08.14.607772

**Authors:** Hong-Dien Phan, Kaitlyn E. Mayne, Willow R.B. Squires, Grant R. Kelly, Reilly H. Smith, Rashid Jafardoust, Sherri L. Christian

## Abstract

CD24 is a glycophosphatidylinositol-linked protein that regulates B cell development. We previously reported that stimulation of CD24 on donor B cells promotes the transfer of functional receptors to recipient B cells via extracellular vesicles (EVs). However, the mechanisms regulating CD24-mediated formation of bioactive EVs are unknown. Using bioinformatics, we found a connection between CD24, and PI3K/AKT and mTOR. To determine if these pathways regulate EV release, we used flow cytometry to follow the transfer of EVs carrying lipid-associated GFP and surface IgM from donor to recipient B cells. Using chemical and genetic inhibition, we found that a PI3K/mTORC2/ROCK/actin pathway regulates bioactive EV formation via activation of acid sphingomyelinase (aSMase) upstream of PI3K. Using single EV analysis, we found that CD24 regulates the formation of the subset of bioactive EVs that are taken up by recipient cells and not total EVs. Interestingly, we also found that ROCK and aSMase modulate ectosome but not exosome formation, when CD24 is stimulated. Lastly, through live cell imaging, we found that PI3K and ROCK are required for inducing membrane dynamics associated with EV formation. These data suggest that this pathway regulates bioactive EV release that, in turn, could regulate B cell development.

**Graphical TOC/Abstract:** 1. Bioinformatics analysis followed by genetic and chemical experiments revealed that a novel pathway regulates EV release downstream of CD24; namely aSMase/PI3K/mTORC2/ROCK/actin.
2. Single EV analysis suggests that the EVs are ectosomes and that CD24 regulates packaging of cargo to create bioactive vesicles rather than regulating total EV release.
3. Live cell imaging shows that CD24 regulates membrane dynamics.

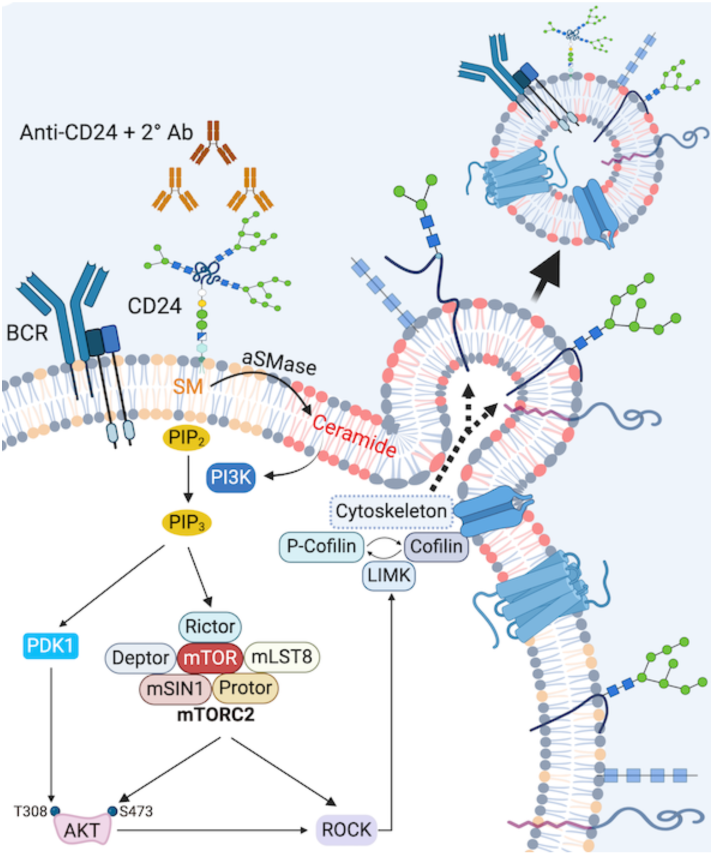

## Introduction

Extracellular vesicles (EVs) are a heterogeneous group of small lipid bilayer-bound particles released by all cells tested to date^1^. There are many kinds of EVs described based on their biogenesis, size, and/or source, including exosomes (30-150nm), and ectosomes (100-1000 nm)^2^. Operationally, EVs are often described based on their size (eg. small vs large EVs) but as exosomes and ectosome overlap in size, their distinguishing feature is actually their biogenesis^3^. Exosomes originate from intraluminal vesicles (ILVs) that form through the inward budding of the endosomal membrane, resulting in the formation of multivesicular bodies (MVB). This is followed by secretion upon fusion of the MVB with the cell plasma membrane. While there are no markers that are exclusively sorted to exosomes vs. ectosomes^2^, exosomes tend to be enriched in a number of proteins including chaperones (Hsp70 and Hsp90), cytoskeletal proteins (actin, myosin, and tubulin), endosomal sorting complex required for transport (ESCRT) proteins (TSG-101 and Alix), proteins involved in vesicle formation and fusion (Rab11, Rab7, Rab2, and LAMP-1), and tetraspanin proteins (CD63, CD81, and CD82)^4–6^. Ectosomes are formed by direct outward budding from the surface of the plasma membrane. Ectosomes tend to be enriched in proteins such as GTP-binding protein, ADP-ribosylation factor 6 (ARF6), matrix metalloproteinases (MMPs), glycoproteins (e.g., GPIb, GPIIb-IIIa), integrins, receptors (e.g., EGFRvIII), other tetraspanin proteins (ie. CD9), Annexin A2, and cytoskeletal elements (e.g., β-actin and α-actinin-4)^7–9^.

Exosome release can be regulated by ESCRT and ESCRT-independent pathways that also participate in ILV formation. For example, release of exosomes is reduced after inhibition of neutral sphingomyelinase (nSMase), a protein responsible for the production of ceramide, suggesting that the budding of ILVs requires ceramide, an important component of lipid raft microdomains^10^. In addition, ApoE and tetraspanin CD63 are recruited for ILV formation and subsequent exosome release without the requirement of ESCRT or ceramide^11, 12^.

Ectosome release can be regulated via several pathways. For example, it has been shown that ectosome release can be stimulated through an increase in intracellular Ca^2+^ concentrations^13^. At steady state, the anionic phospholipids, phosphatidylserine (PS) and phosphatidylethanolamine (PE) localize to the inner leaflet of the plasma membrane, while phosphatidylcholine (PC) and sphingomyelin (SM) are found on the external membrane leaflet^14^. The physiological membrane asymmetry is maintained by five transmembrane enzymes: gelsolin, scramblase, flippase, translocase, and calpain^15^. The increase in cytoplasmic Ca^2+^ levels inhibits the aminophospholipid translocase and activates lipid scramblase simultaneously^16, 17^. This process drives rearrangements in the asymmetry of the membrane phospholipids to expose PS from the inner leaflet to the cell surface. It then leads to the physical collapse of cell membrane asymmetry, which can promote ectosome release^18^.

In addition to lipids, cytoskeletal elements and their regulators are required for ectosome formation^19^. Reducing the levels of phosphatidylinositol 4,5-bisphosphate (PI(4,5)P_2_), which participates in anchoring the membrane to the cortical cytoskeleton, can induce the disruption of the cortical cytoskeleton interaction with the plasma membrane^20, 21^, leading to increased ectosome biogenesis^22^. Reduction of PI(4,5)P_2_ can occur via activation of phospholipase C (PLC)-γ, phospholipase D (PLD), phosphoinositide 3-kinases (PI3Ks), or phosphatidylinositol phosphatases^23^. Rab22a and ARF6, members of the Ras GTPase family, also have essential roles in ectosome generation. Rab22A is directly involved in ectosome formation as evidenced by the fact that knockdown prevents vesicle release^24^. Importantly, ARF6 activity is required for subsequent phospholipase D activation, leading to localized myosin light chain kinase activity at the neck of budding vesicles^25^. Furthermore, ARF6-mediated activation of RhoA and Rho-associated kinase (ROCK) signaling has been implicated in ectosome formation^25, 26^. An additional study has shown that the antagonistic interaction between Rab35 and ARF6 also controls ectosome biogenesis^27^. Furthermore, a key regulator in the formation of ectosomes, Ca^2+^ ions also contribute to the reorganization of the cytoskeleton through the activation of cytosolic calpain protease^28^. Calpain cleaves several cytoskeletal components such as actin, ankyrin, protein 4.1, and spectrin^29, 30^. Calpain-mediated cleavage of the cytoskeleton further disrupts the cortical cytoskeleton protein network, consequently, allowing membrane budding^31^. Lastly, similar to exosomes, ceramide production can increase ectosome release^32^.

CD24 (also known as heat stable antigen) is a glycophosphatidylinositol (GPI)-anchored glycoprotein, which is localized to lipid rafts on the plasma membrane^33^. It contains 27 amino acids with extensive N-linked and O-linked glycosylation that can result in variable molecular weight products. CD24 is expressed on several cell types, including B cells, T cells, neutrophils, eosinophils, dendritic cells, macrophages, epithelial cells, and cancer cells. During B cell development in the bone marrow, CD24 is first expressed by pro-B cells (also called Fraction B) but is also highly expressed at the pre-B cell (Fraction C, C’, and D) stages and in transitional B cells^34^.

One of the most well-described effects of CD24-mediated signaling is its promotion of apoptosis in developing B cells^35^. CD24 recruits Src family protein tyrosine kinases (PTKs) to activate signalling pathways via direct protein phosphorylation, intracellular calcium mobilization, and transcription factor activation^34^. Moreover, existing evidence has demonstrated that multiple cancer related signaling pathways, such as Wnt/β-catenin, mitogen activated protein kinase (MAPK), Src or PI3K/Akt, Notch, and Hedgehog, are activated downstream of CD24^36^.

We previously discovered that the engagement of CD24 enhances the release of EVs from *ex vivo* bone marrow-derived B cells and the mouse WEHI-231 B cell lymphoma cell lines^37^. We also found that the RNA cargo and the EV proteome are relatively stable, but the composition of the membrane proteins on the EVs is altered^38^. Recently, we employed a model system where donor cells expressing palmitoylated GFP (WEHI-231-GFP cells) were co-cultured, after CD24 stimulation, with recipient cells lacking IgM and expressing palmitoylated tdTomato (WEHI-303-tdTomato cells) to study EV-mediated transfer of functional receptors^39^. We found that EVs trafficked lipid and membrane proteins between B lymphocytes in response to stimulation of CD24 on the donor cells. Notably, the transported receptors can induce apoptosis, which may affect B cell development in recipient bystander B cells during B cell development^39^. However, the underlying mechanisms that govern EV formation and subsequent uptake in recipient cells in response to engagement of CD24 have not been described.

To better understand the pathways that may regulate EV transfer in response to CD24, we first analyzed which genes were differentially expressed in the same manner as CD24 across B cell development using a dataset from the Immunological Genome Project (ImmGen) database^40^. Network analysis revealed that CD24 expression is associated with genes that are enriched in the PI3K/Akt/mTOR signaling pathway. PI3K/Akt signaling is implicated in various cellular processes, including differentiation, growth, proliferation, and intracellular trafficking, all of which are factors in B cell development and activation^41, 42^. mTOR is found in two complexes: mTORC1 (which contains mTOR, Raptor, and other proteins) and mTORC2 (which contains mTOR, Rictor, and other proteins)^43^. mTORC1, called the rapamycin-sensitive mTOR complex, is a key regulator of cell proliferation and survival downstream of the PI3K/Akt pathway. mTORC2 triggers Akt activation by phosphorylation on serine 473, as well as regulation of the actin cytoskeleton via RhoA and PKCα^44^. Regulation of phosphoinositides by PI3K and the actin cytoskeleton by RhoA can regulate ectosome release^20, 45^. Therefore, we addressed the hypothesis that CD24-mediated EV transfer is regulated by the PI3K/Akt/mTOR pathway. Using our co-culture model, we found that CD24 mediates EV transfer via the activation of acid sphingomyelinase (aSMase), which activates the PI3K/mTORC2/ROCK pathways followed by actin cytoskeletal rearrangement. Interestingly, total EV release was not affected by CD24 stimulation suggesting that CD24 regulates formation of bioactive EVs via packaging of specific cargo. Lastly, live image analysis revealed that PI3K and ROCK regulate membrane dynamics associated with EV release.

## Materials and Methods

### Cell culture

WEHI-231 cells (American Type Culture Collection (ATCC), Manassas, VA) and WEHI-303.1.5 (WEHI-303)^46^ were transfected with a lentiviral plasmid encoding either palm-GFP (WEHI-231-GFP) or palm-tdTomato (WEHI-303-tdTomato) obtained from Charles Lai, Institute of Atomic and Molecular Sciences, Taiwan^47^. Cells were cultured in RPMI-1640 medium (Gibco) containing 10% heat inactivated fetal bovine serum (FBS) (Gibco), 1.0 mM sodium pyruvate (Gibco), 50 mM 2-mercaptoethanol (Sigma), and 100 U/ml penicillin, 100 µg/ml streptomycin (Invitrogen). For knockdown experiments, WEHI-231-GFP cells (2 x 10^5^) were transiently transfected with 2 mM siRNA with the Neon Transfection Kit (MPK1025B, Invitrogen) using the Neon Electroporation system according to the manufacturer’s instructions. The siRNAs used were control non-targeting siRNA-A (Santa Cruz Biotechnology, SC-37007), PI 3-kinase p110δ siRNA (Santa Cruz Biotechnology, SC-39132), and Rock-1 siRNA (Santa Cruz Biotechnology, SC-36432). Cells were pulsed once with a voltage of 1700 and a width of 20. Transfected cells were cultured for 55 h before use.

### Bioinformatics

ImmGen data (GEO accession number GSE15907), containing gene expression data for B cells in the Fraction (Fr) A-F stages of development, were used for this analysis. The microarray gene expression data files for Fr A, Fr B/C, Fr C’, Fr D, and Fr F were background corrected, and robust multi-array average (RMA) normalized using the Oligo, Biobase, and pd.mogene.1.0.st.v1 Bioconductor packages in R version 4.0.0. A list of differentially expressed genes with similar expression patterns to *Cd24a* was then compiled using the Limma and Affycoretools Bioconductor packages. Fr A B cells (pre-pro B cells) was used as the negative control for the differential expression analysis. The gene transcript cluster ID list was annotated with the corresponding gene names using the NetAffx analysis batch query and merged with the R expression data to create a median linkage hierarchical cluster in Genesis version 1.8.1. Pathway networks were generated in Cytoscape version 3.8.0 using the ClueGO plugin version 2.5.7. The analysis used pathway network data from the KEGG, REACTOME Pathways, and WikiPathways databases.

### Phosphatidylinositol (3,4,5)-trisphosphate (PIP_3_) Quantification

WEHI-231-GFP cells (0.5 x 10^6^) were pre-treated for 15 min with the PI3K inhibitor, 10µM LY294002 (9901, Cell Signaling Technology), 10 µM imipramine (J63723.06, Alfa Aesar), or with an equal volume of DMSO. Cells were then stimulated with 10 µg/ml of functional grade primary monoclonal M1/69 rat anti-mouse CD24 antibody (16-0242085, eBioscience) or 10 µg/ml matching primary isotype antibody (16-4031-85, eBioscience) that was pre-incubated with 5 µg/ml goat anti-rat secondary antibody (112-005-003, Jackson ImmunoResearch) for 15 min.

After treatment, cells were centrifuged and phosphoinositides were extracted and measured using the PIP_3_ ELISA kit (Creative Diagnostic, DEIA-XYZ6). All experiments were performed at least four times, each carried out in duplicate according to the manufacturer’s instructions.

### Analysis of cell surface IgM and lipid transfer by flow cytometry

WEHI-231-GFP cells (0.5 x 10^6^) were pre-treated with either DMSO or inhibitor for 15 min. The following inhibitors were used: 10 µM LY294002, 0.25 µM MK2206 (S1078, Selleckchem), 0.25 µM Torin 1 (14379S, Cell Signaling Technology), 0.1 µM Rapamycin (A8167, APExBIO), 0.1 µM JR-AB2-011 (HY-122022, MedChemExpress), 10 µM Y27632 (13624S, Cell Signaling Technology), 0.2 µM Cytochalasin D (PHZ1063, Gibco, ThermoFisher), 10 µM imipramine, 20 µM GW4869 (D1692, Sigma Aldrich), 20 µM ARC39 (13583, Cayman Chemical), or 0.5 µg/mL Ionomycin (I24222, ThermoFisher). Cells were stimulated with anti-mouse CD24 antibody or isotype antibody as above followed by centrifugation to remove excess antibody. Cells were then co-cultured with the recipient WEHI-303-tdTomato cells (0.5 x 10^6^) for 24 h. To assess cell surface IgM and lipid transfer, the mixed cells were resuspended in ice-cold FACS buffer (1x PBS, pH 7.4, containing 1% heat-inactivated FBS) and then stained on ice for 30 min with 0.5 mg of anti-mouse IgM-PE-Cy7 (25-5890, eBioscience) to detect IgM. Cells were then washed with FACS buffer and analyzed by flow cytometry. Flow cytometry was performed using a CytoFLEX (Beckman Coulter) counting at least 10,000 events and data were analyzed using FlowJo software version 10.4.1. The experimental design as well as gating and analysis strategies can be found in Figure S1-2.

### Immunoblotting

Cell extracts were separated on 8% to 15% SDS-PAGE gels and transferred to nitrocellulose membranes, which were blocked with either 5% milk powder or 5% BSA in 0.1% Tween-20-Tris-buffered saline (TBST) for 1 h at room temperature. The membranes were incubated with primary antibody in TBST overnight at 4°C: Phospho-Akt (Ser473) antibody (Cell Signaling Technology, #9271, 1:1000), Akt (Cell Signaling Technology, #9272, 1:1000), Phospho-cofilin (Cell Signaling Technology, #3313; 1:1000), cofilin (Cell Signaling Technology, #3318, 1:1000), Rock-1 antibody (G-6) (Santa Cruz Biotechnology, SC-17794, 1:500), PI3K p110δ antibody (A-8) (Santa Cruz Biotechnology, SC-55589, 1:1000), and GAPDH (G-9) (Santa Cruz Biotechnology, SC-365062, 1:1000). The membrane was washed and incubated with a horseradish peroxidase (HRP)-conjugated goat anti-rabbit IgG (H+L) (Bio Rad, #1706515) or goat anti-mouse IgG (H+L) (Bio Rad, #1721011). Immunoreactive bands were visualized on a Chemidoc gel system (BioRad) using an ECL substrate (WBULS0100, Millipore). Image manipulation involved adjustments to brightness and contrast only.

### Analysis of EV surface markers and size by flow cytometry

WEHI-231-GFP cells (0.5 x 10^6^) were pre-treated with inhibitors (LY294002, Y27632, imipramine, ARC39) and stimulated for 15 min as above. They were then centrifuged at 500 x g for 5 mins to remove excess antibody and resuspended in EV-depleted media. The cells were then incubated for 1 h at 37°C in 5% CO_2_ followed by centrifugation 500xg for 5 min. The EV supernatant was further clarified by centrifugation at 2000 x g for 5 min. EV-depleted medium was prepared according to the previously established protocol^48^; briefly, complete medium with 20% FBS was centrifuged at 100,000 × g at 4 C for 16 h, passed through a 0.22-mm filter, and then mixed with serum-free RPMI 1640 in a 1:1 ratio. EVs were diluted according to the Cellarcus Bioscience CytoFLEX Protocols for vFC™ vesicle flow cytometry EV analysis assay (CBS4-1, Cellarcus Biosciences, La Jolla, California). We determined the optimal dilution of EVs was 1:400 in vFC Staining Buffer. EVs were stained with vFRed membrane stain according to manufacturer’s protocols and the following antibodies: 1μg anti-Annexin-2/ANXA2 Antibody (C-10) Alexa Fluor 647 (Sc-28385, Santa Cruz), 1μg anti-CD9 (KMC8) eFluor 450 (48-0091-82, ThermoFisher), 1μg anti-CD63 (NVG-2) PE (12-0631-82, ThermoFisher), and 1μg anti-CD107a (LAMP-1) Antibody (1D4B) PE-Cy7 (25-1071-82, ThermoFisher) for 1 h in the dark. The following isotype staining controls were used at 1 μg per sample, IgG2a kappa isotype control Alexa Fluor 647 (51-4724-81, ThermoFisher), Rat IgG2a kappa Isotype Control (eBR2a) PE (12-4321-80, ThermoFisher), Rat IgG2a kappa isotype control (eBR2a) eFluor 450 (48-4321-82, ThermoFisher), and Rat IgG2a kappa Isotype Control (eBR2a), PE-Cyanine7, eBioscience (25-4321-82, ThermoFisher). Flow cytometry was performed using a CytoFLEX (Beckman-Coulter). Custom parameters were set according to Cellarcus Biosciences vFC Instrument Setup for Beckman-Coulter CytoFLEX and optimized using manufacturer’s protocols with vCal^TM^ nanoRainbow Beads (CBS6-2.5ml), vCal^TM^ anti-mouse nanoCal^TM^ beads (CBS7M-2.5ml), and vCal^TM^ anti-rat nanoCal^TM^ beads (CBS7RT-2.5ml). Stained vesicles were detected using a vFRed trigger threshold manually set to 1900 based on background from free dye in buffer. Data was collected by running samples at 60 μL/min for 120 s on the CytoFLEX. Data was analyzed using vFC^TM^ Data Analysis FCS Express Analysis Layouts for the CytoFLEX and FCS Express (De Novo Software version 7.22.0031, Pasadena, California). Experiments were performed four times and gated by a collection time of 15 s-70 s.

### Live cell imaging

Glass coverslips (35 mm) were cleaned thrice with 1x PBS and incubated with 1 mg/mL of anti-MHC class II antibody (Sigma, MABF33) overnight at 4°C. Then, 0.5 x 10^6^ WEHI-231-GFP cells were seeded onto the glass coverslips for 30 min at 4°C. The cells were washed with PBS and visualized with a Carl Zeiss LSM 880 laser scanning microscope system with a 63x oil immersion lens with numerical aperture (NA) of 1.4 objective under environment at 37^°^C and 5% CO_2_. Laser excitation light was provided at a wavelength of 488 nm, and fluorescent emissions were collected at wavelengths above 515 nm. For image acquisition, an exposure time of 0.8 second was adopted with a binning of 2 x 2 yielding a pixel size of 0.68 mm. Individual cells within a single field of view were captured with an Airyscan detector over a 5-min period for 250 cycles, with a 1.2-second shuttered interval between each image. Raw Airyscan images were processed using the Zeiss Zen Blue analysis software. Subsequent image processing was conducted in ImageJ to create a video with 25 frames per second for a 10-second length.

Analysis of videos was done in ImageJ using the analyze_blebs plugin^49^ to determine the coefficient of variance of the area of individual cells over a time series of 10 image spanning the 5-min period.

### Statistical analysis

One-way ANOVA and two-way ANOVA with Sidak’s post-hoc test were performed using GraphPad Prism (version 10.1.1) software, as indicated. Data were confirmed to be parametric based on QQ plots. Live cell imaging data was graphed and analyzed in R using ggplot2^50^ and baseR, respectively.

## Results

Prior research from our lab showed that B cells release EVs in response to CD24 engagement^37–39^. As the first step in attempting to elucidate the pathway(s) responsible for CD24-mediated release of EVs, we performed a bioinformatics-based analysis to find potential signaling pathways linked to CD24 expression in developing B cells. Differential expression analysis produced a list of 1838 genes expressed in a similar manner as *Cd24a*. We identified 44 genes in the clade that most closely resembled *Cd24a* (Figure 1A). Of these, 41 could be used to predict CD24-related signaling pathways. In the networks of connected pathway terms, seven of the nodes had terms related to PI3K/Akt/mTOR signalling (Figure 1B). Thus, we predicted that CD24 may activate PI3K signaling. To determine if engagement of CD24 activated PI3K, we analyzed the levels of PIP_3,_ the immediate product of PI3K activation. We found that PIP_3_ levels were significantly increased in response to CD24 stimulation, but not with isotype-control stimulated cells. In addition, we observed the increase of PIP_3_ was inhibited in the presence of the PI3K inhibitor, LY294002 (LY, Figure 1C). These results clearly demonstrate that CD24 triggers B cells to activate PI3K.

**Figure 1.**
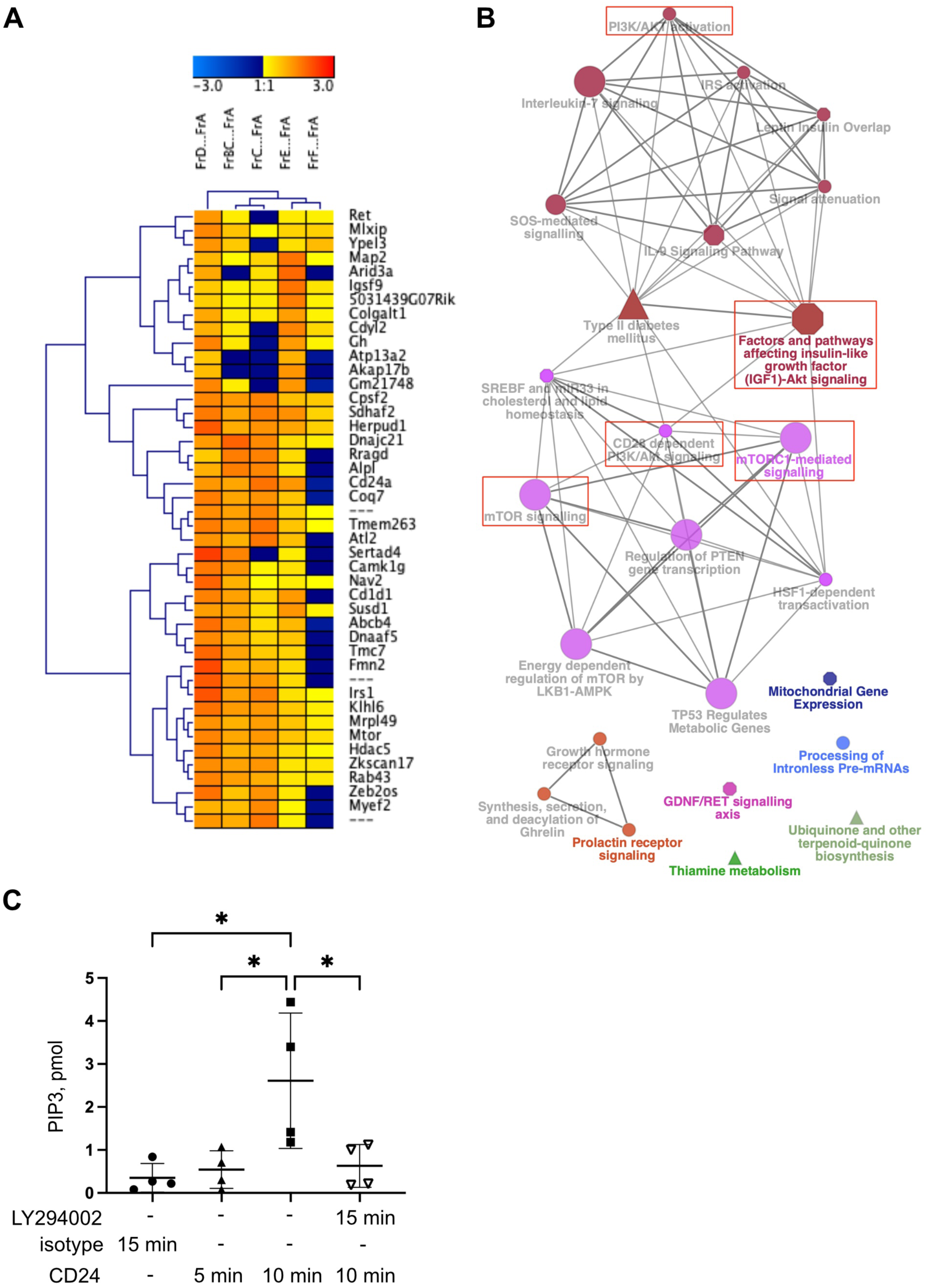
CD24 expression is associated with the PI3K-Akt signaling and mTOR pathways. (A) Hierarchical cluster analysis showing the clade of 44 genes (41 annotated) with similar differential gene expression patterns as *CD24a*. (B) Pathway network analysis generated from the differentially expressed 41-gene list identified the PI3K/Akt and mTOR signaling pathway as being associated with CD24 expression in developing B cells. The red boxes indicate nodes with terms related to PI3K-Akt or mTOR signaling. (C) WEHI-231-GFP cells were pre-treated with LY294002 or DMSO (vehicle control) for 15 min then cells were stimulated with isotype antibody (isotype) or anti-CD24 stimulating (CD24) antibody for the times indicated. PIP_3_ levels were analyzed by ELISA, n=4, significance was determined by a one-way ANOVA followed by the Sidak’s multiple comparison test *P<0.05.

To determine if the PI3K signaling pathway regulates EV transfer in response to CD24 stimulation, we used our model system to track EV transfer (Supplemental Figure S1A-C). Transfer of EVs, as shown previously^39^, can be evaluated by the uptake of GFP and IgM in the recipient tdTomato-positive cells. Gating on live cells gave the same results as a more narrow gating on single cells, giving us confidence that we are analyzing cells that have taken up EVs and not doublets (Supplemental Figures S1 and S2). As we have previously shown, we found a significant increase in lipid transfer between the CD24-stimulated and the isotype-stimulated group (Figure 2A). The amount of lipid transferred in response to CD24 was significantly reduced in the presence of LY (Figure 2A and Supplemental Figure S2A and C). Parallel investigations into protein transfer (IgM) revealed a similar pattern. There was a significant increase in protein transfer between the CD24-stimulated and the isotype-stimulated group in the presence or absence of LY with significantly less transfer in the CD24-stimulated group in the presence of LY (Figure 2B and Supplemental Figure S2B and D). To verify inhibition of the PI3K pathway, we analyzed Akt phosphorylation. We found that the level of phosphorylated AKT increased in response to CD24 stimulation, while it was markedly lower in cells pre-treated with LY compared with those that received DMSO (Figure 2C). Thus, these data show that LY inhibits CD24-mediated PI3K signaling; however, some residual activity remains.

**Figure 2.**
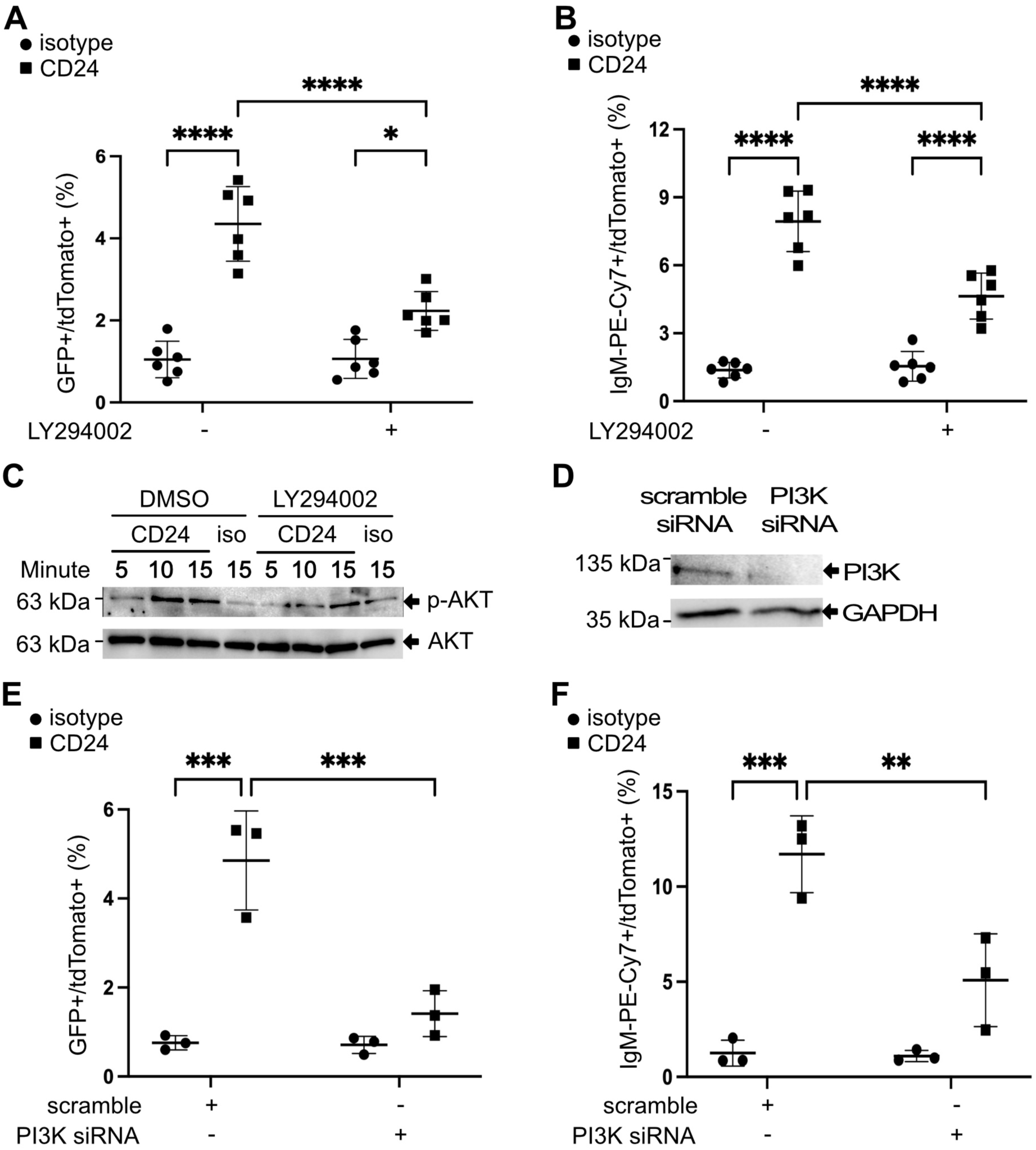
CD24-mediated EV transfer is regulated by PI3K. (A-B) WEHI-231-GFP cells were pre-treated with LY294002 or DMSO for 15 min, then stimulated with anti-CD24 (CD24) or isotype control (isotype, iso) for 15 min, followed by washout and then a 24 h co-culture with WEHI-303-tdTomato cells. (A) Percent GFP and tdTomato double-positive cells and (B) Percent IgM and tdTomato double-positive cells after 24 h incubation. n=6, statistical significance determined by a two-way ANOVA (interaction significant at P=0.0003 for A and P=0.0002 for B) followed by the Sidak’s multiple comparison test *P<0.05, ****P<0.001. C) Total cell lysates from WEHI-231-GFP cells pre-treated with DMSO or LY2940002 for 15 min, then stimulated with the above antibodies for different times indicated. Phosphorylated Akt and total Akt expression levels were determined by immunoblotting. D) WEHI-231-GFP cells were transfected with scrambled control siRNA or PI3K siRNA, and PI3K and GAPDH levels determined by immunoblotting. Shown are representative western blots from 3 replicates with molecular weight marker positions indicated on the left of each blot. (E-F) WEHI-231-GFP cells with or without PI3K siRNA knock-down were stimulated with anti-CD24 or isotype for 15 min, followed by washout and then a 24 h co-culture with WEHI-303-tdTomato cells. (E) Percent GFP and tdTomato double-positive cells and (F) Percent IgM and tdTomato double-positive cells after 24 h incubation. n=3, statistical significance determined by a two-way ANOVA (interaction significant at P=0.0015 for E and P=0.0088 for F) followed by the Sidak’s multiple comparison test **P<0.01, ***P<0.005.

Next, we sought to genetically validate the LY result using siRNA knock-down of PI3K in donor cells. The absence of PI3K expression in siRNA-transfected WEHI-231-GFP cells was confirmed by Western blot (Figure 2D). Analysis of lipid and protein transfer between the CD24-stimulated and isotype-stimulated groups revealed a significant increase in the scrambled control siRNA group, as expected. However, there was no transfer of lipid or protein in either the isotype-stimulated or CD24-stimulated groups when PI3K was knocked-down in the donor cells (Figure 2E-F). These findings strongly support a role for the PI3K signaling pathway in regulating CD24-mediated production of bioactive EVs that are taken up by recipient cells.

We investigated further to see if proteins downstream of PI3K are involved in EV transfer in response to CD24. After pre-treatment with the Akt inhibitor MK-2206, we found a significant decrease in CD24-mediated lipid and protein transfer to recipient cells (Figure 3A-B). Similar to PI3K inhibition, some residual EV transfer remained in the presence of the inhibitor.

**Figure 3.**
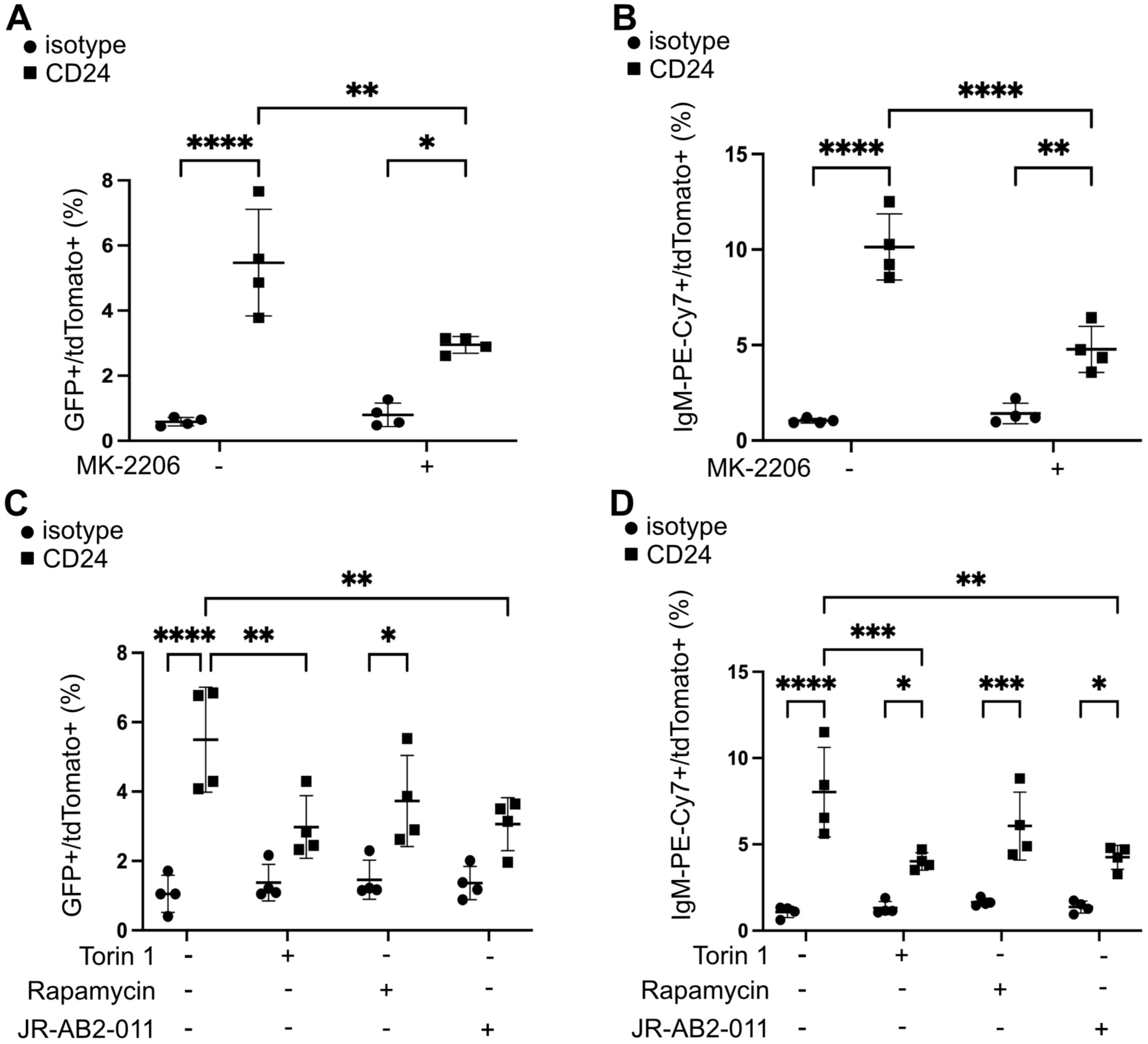
CD24-mediated EV transfer is dependent on Akt and mTOR signaling pathways. (A-B) WEHI-231-GFP cells were pre-treated with MK-2206 or DMSO for 15 min, then stimulated with anti-CD24 (CD24) or isotype control (Isotype) for 15 min, followed by washout and then a 24 h co-culture with WEHI-303-tdTomato cells. (A) Percent GFP and tdTomato double-positive cells and (B) Percent IgM and tdTomato double-positive cells after 24 h incubation. n=4, statistical significance determined by a two-way ANOVA (interaction significant at P=0.0074 for A and P=0.0002 for B) followed by the Sidak’s multiple comparison test *P<0.05, **P<0.01, ****P<0.001. (C-D) WEHI-231-GFP cells were pre-treated with Torin 1 or Rapamycin or JR-AB2-011 or DMSO for 15 min, then stimulated with anti-CD24 or isotype control for 15 min, followed by washout and then a 24 h co-culture with WEHI-303-tdTomato cells. (C) Percent GFP and tdTomato double-positive cells and (D) Percent IgM and tdTomato double-positive cells after 24 h incubation. n=4, statistical significance determined by a two-way ANOVA (interaction significant at P=0.0143 for C and P=0.0066 for D) followed by the Sidak’s multiple comparison test *P<0.05, **P<0.01, ***P<0.005, ****P<0.001.

Other signalling pathways downstream of PI3K are the mTOR signalling pathways. mTOR forms two structurally and functionally distinct complexes: mTORC1 and mTORC2^51^ Therefore, we next determined if mTORC1 or mTORC2 regulates CD24-mediated EV release. To do this, we used Torin 1, an inhibitor of mTORC1/2, rapamycin, an inhibitor of mTORC1, and JR-AB2-011, an inhibitor of mTORC2^52^. As can be seen in Figure 3C and Figure 3D, there was a substantial statistical difference in lipid and protein transfer between the mTORC2 inhibition groups (Torin 1 and JR-AB2-011) and the control stimulated group, indicating that mTORC2 controls EV transfer in response to CD24.

Rho-associated protein kinase (ROCK) is known to be regulated by the PI3K/mTORC2 signaling pathway^53^ and was identified by the bioinformatics analysis (Figure 1B). Therefore, we next asked whether CD24-mediated EV transfer was similarly regulated by the ROCK signaling pathway using the inhibitor Y27632^54^. We found that, in comparison to EVs from the control group, the release of EVs from donor cells pre-treated with Y27632 resulted in a significantly lower transfer of lipid and protein to recipient cells upon CD24 stimulation (Figure 4A-B). We found that there was a reduction but not a total block in phosphorylated cofilin, a downstream target of ROCK, suggesting that the inhibitor did not fully block ROCK activity (Figure 4C).

**Figure 4.**
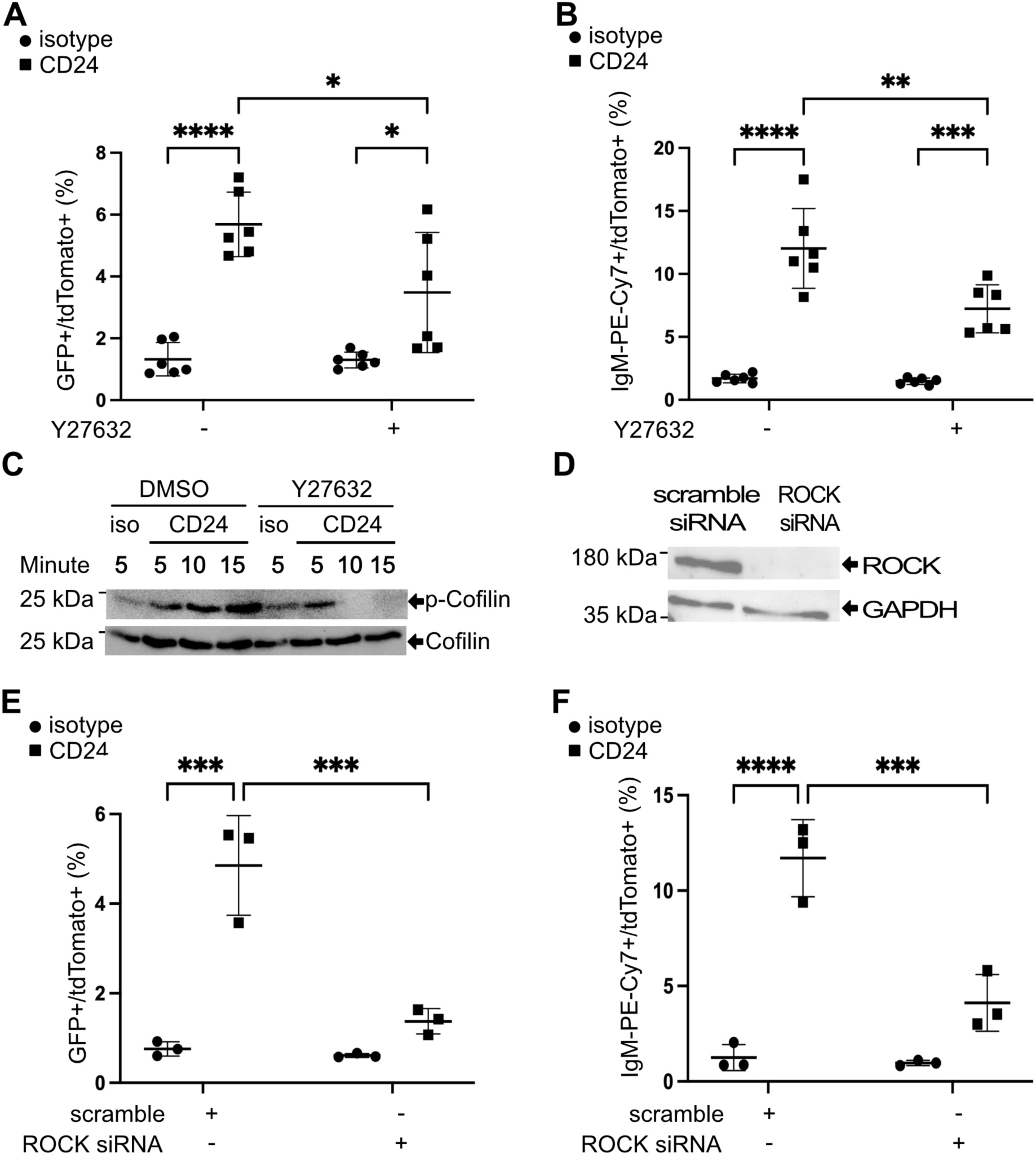
CD24-mediated EV transfer is dependent on ROCK. (A-B) WEHI-231-GFP cells were pre-treated with Y27632 or DMSO for 15 min, then stimulated with anti-CD24 (CD24) or isotype control (isotype, iso) for 15 min, followed by washout and then a 24 h co-culture with WEHI-303-tdTomato cells. (A) Percent GFP and tdTomato double-positive cells and (B) Percent IgM and tdTomato double-positive cells after 24 h incubation. n=6, statistical significance determined by a two-way ANOVA (interaction significant at P=0.03 for A and P=0.0069 for B) followed by the Sidak’s multiple comparison test *P<0.05, **P<0.01, ***P<0.005, ****P<0.001. C) Total cell lysates from WEHI-231-GFP cells pre-treated with DMSO or Y27632 for 15 min, then stimulated with the above antibodies for different times indicated. Phosphorylated Cofilin and total Cofilin expression levels were determined by immunoblotting. Shown are representative western blots from 3 replicates with molecular weight marker positions indicated on the left of each blot. D) WEHI-231-GFP cells were transfected with scrambled control siRNA or ROCK siRNA, and ROCK and GAPDH levels determined by immunoblotting. (E-F) WEHI-231-GFP cells with or without ROCK siRNA knock-down were stimulated with anti-CD24 or isotype for 15 min, followed by washout and then a 24 h co-culture with WEHI-303-tdTomato cells. (E) Percent GFP and tdTomato double-positive cells and (F) Percent IgM and tdTomato double-positive cells after 24 h incubation. n=3, statistical significance determined by a two-way ANOVA (interaction significant at P=0.0011 for E and P=0.0013 for F) followed by the Sidak’s multiple comparison test ***P<0.005, ****P<0.001.

Next, to genetically validate the Y27632 inhibitor result, we used siRNA to knock-down ROCK in donor cells (Figure 4D). We found that there was no transfer of lipids or proteins with CD24 stimulation when ROCK was knocked-down (Figure 4E-F). These data clearly demonstrate that CD24-mediated EV generation is regulated by ROCK.

We hypothesized that dynamic cytoskeletal reorganization downstream of ROCK that changes the shape of cell membrane will, in turn, promote bioactive EV formation and subsequent uptake. Donor cells were thus pre-treated with cytochalasin D to prevent cytoskeleton rearrangement.

The results showed that there was no increase in lipid and protein transfer in the cytochalasin D-treated CD24-stimulated group compared to the control stimulation (Figure 5A-B). Thus, actin cytoskeleton regulation plays a key role in regulating CD24-mediated EV transfer.

**Figure 5.**
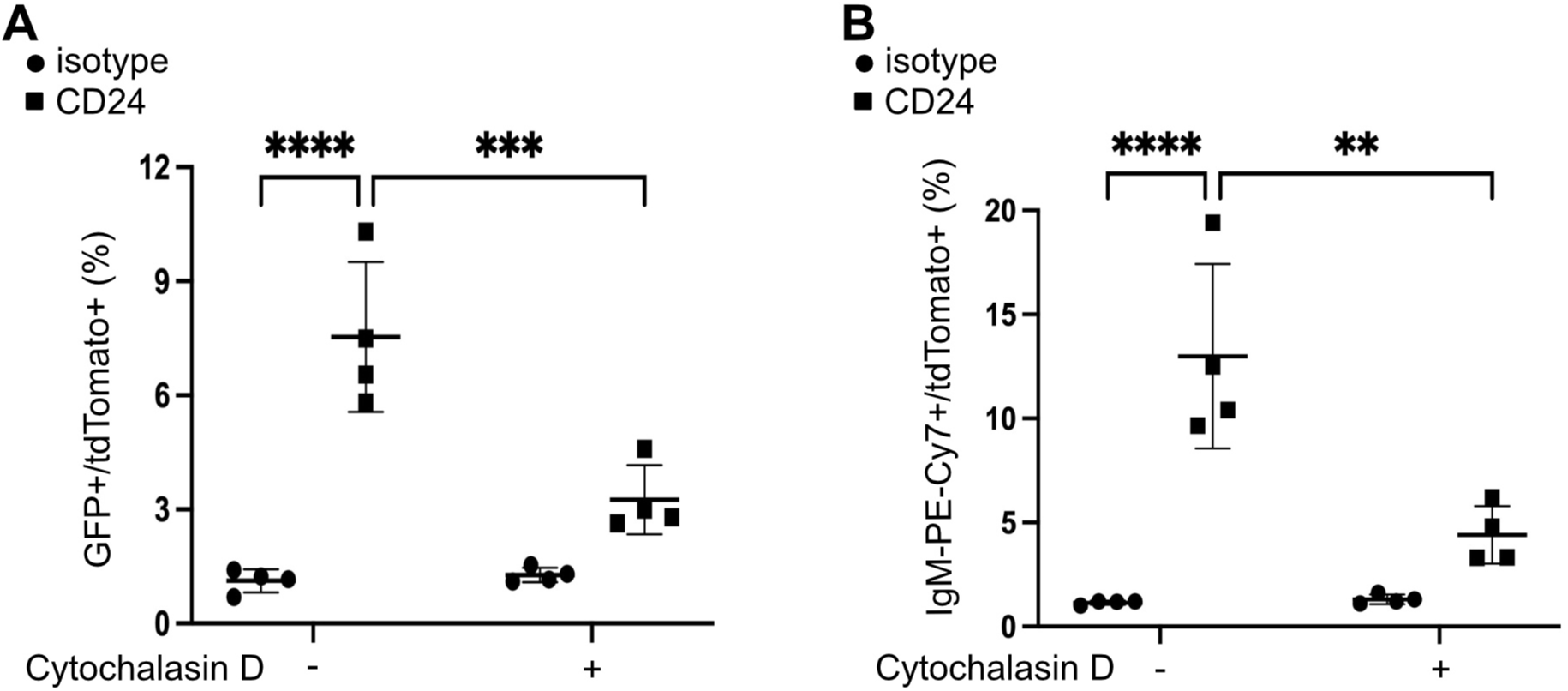
CD24-mediated EV transfer is controlled by actin cytoskeleton re-organization. (A-B) WEHI-231-GFP cells were pre-treated with Cytochalasin D or DMSO for 15 min, then stimulated with anti-CD24 (CD24) or isotype control (isotype) for 15 min, followed by washout and then a 24 h co-culture with WEHI-303-tdTomato cells. (A) Percent GFP and tdTomato double-positive cells and (B) Percent IgM and tdTomato double-positive cells after 24 h incubation. n=4, statistical significance determined by a two-way ANOVA (interaction significant at P=0.0017 for A and P=0.0027 for B) followed by the Sidak’s multiple comparison test **P<0.01, ***P<0.005, ****P<0.001.

**Figure 6.**
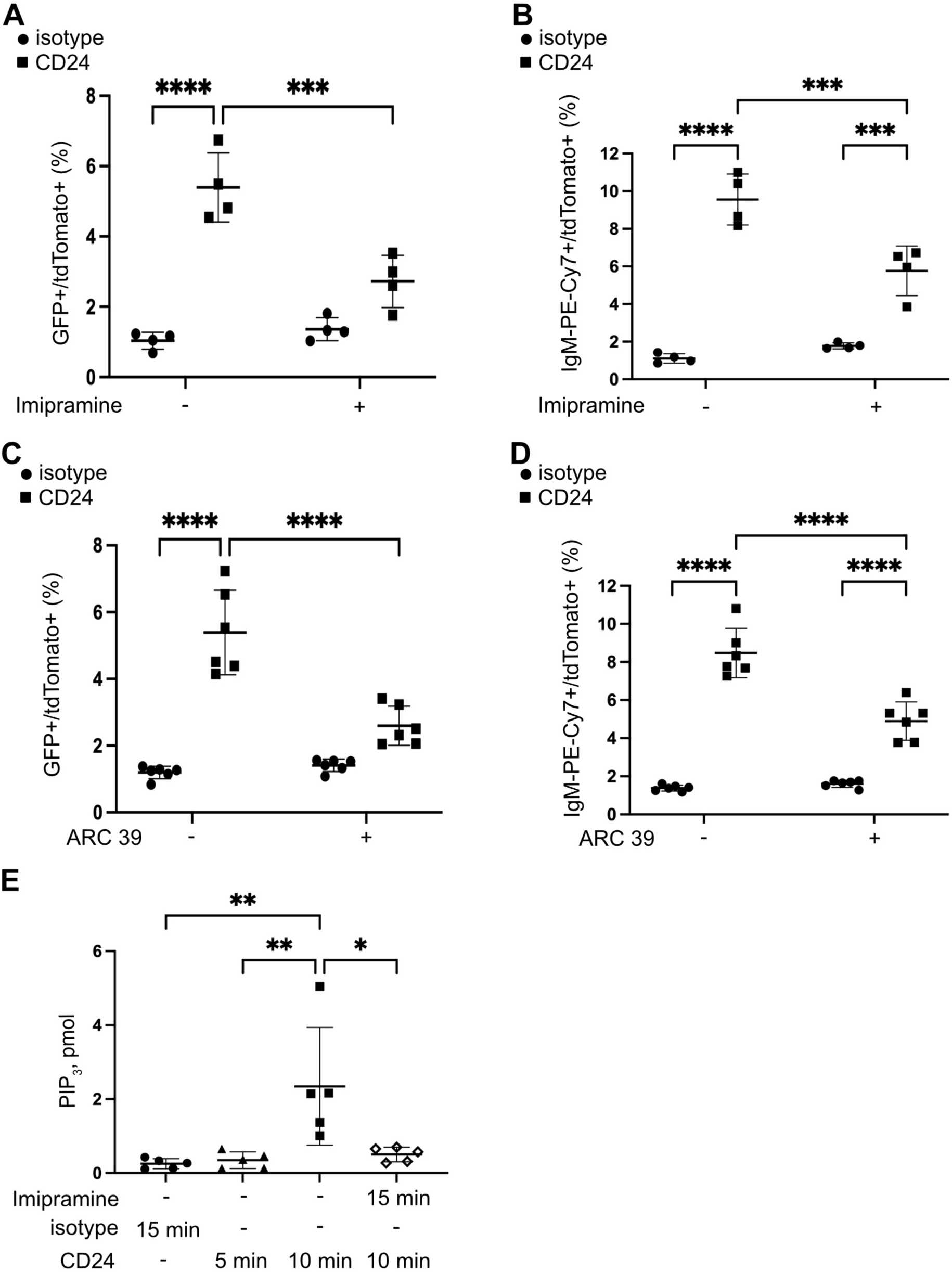
CD24-mediated EV transfer is controlled by aSMase activity. (A-B) WEHI-231-GFP cells were pre-treated with imipramine or DMSO for 15 min, then stimulated with anti-CD24 (CD24) or isotype control (isotype) for 15 min, followed by washout and then a 24 h co-culture with WEHI-303-tdTomato cells. (A) Percent GFP and tdTomato double-positive cells and (B) Percent IgM and tdTomato double-positive cells after 24 h incubation. n=4, statistical significance determined by a two-way ANOVA (interaction significant at P=0.0006 for A and P=0.0006 for B) followed by the Sidak’s multiple comparison test ***P<0.005, ****P<0.001. (C-D) WEHI-231-GFP cells were pre-treated with ARC39 or DMSO for 15 min, then stimulated with anti-CD24 (CD24) or isotype control (isotype) for 15 min, followed by washout and then a 24 h co-culture with WEHI-303-tdTomato cells. (C) Percent GFP and tdTomato double-positive cells and (D) Percent IgM and tdTomato double-positive cells after 24 h incubation. n=6, statistical significance determined by a two-way ANOVA (interaction significant at P<0.0001 for C and P<0.0001 for B) followed by the Sidak’s multiple comparison test ***P<0.005, ****P<0.001. (E) WEHI-231-GFP cells were pre-treated with imipramine or DMSO (vehicle control) for 15 min then cells were stimulated with isotype antibody (isotype) or anti-CD24 stimulating antibody (CD24) for the times indicated. PIP_3_ levels were analyzed by ELISA, n=5, significance was determined by a one-way ANOVA followed by the Sidak’s multiple comparison test *P<0.05, **P<0.01.

Sphingomyelin (SM), a phospholipid that localizes on the plasma membrane’s outer leaflet and has a strong affinity for cholesterol, plays a significant role in determining the fluidity and structural integrity of the plasma membrane^55^. nSMase and aSMase catalyze the hydrolysis of SM to ceramide, a process that is involved in both exosome and ectosome release^56^, although the involvement of nSMase has recently been challenged^57^. nSMase activity can be blocked by GW4869^12^, while a previous study has shown that imipramine blocked ectosome formation from glial cells by inhibiting aSMase^32^. Thus, we treated donor cells with either GW4869, imipramine or the more specific aSMase inhibitor ARC39 to determine if nSMase or aSMase regulates CD24-mediated EV transfer. We found that there was no difference in lipid and protein transfer with GW4896 (Supplemental Figure 2A-B). In contrast, CD24-mediated lipid and protein transfer were significantly inhibited with imipramine or ARC39 treatment (Figure 5C-D). Thus, aSMase, but not nSMase, regulates CD24-mediated EV transfer.

We next determined if PI3K was upstream or downstream of aSMase by determining if inhibition of aSMase alters PIP_3_ levels in response to CD24 stimulation. We found that the CD24-mediated increase of PIP_3_ was significantly inhibited in the presence of imipramine (Figure 5E). Therefore, we conclude that aSMase is upstream of PI3K.

The altered uptake seen in recipient cells could be due to either alteration of EV release or changes to EV cargo regulating uptake. Therefore, we directly analyzed single EVs using flow cytometry to enumerate EVs released by donor cells. We also co-stained for markers known to be enriched in exosomes compared to ectosomes. We chose CD63 and LAMP-1 as exosomal markers, and CD9 and Annexin A2 (ANXA2) as ectosome markers, based on previously reported biomarker expression^6, 9, 58, 59^. Similar to our previous work^37, 39^, we found that CD24 stimulation did not significantly change the number of total (Figure 7A) or GFP-positive EVs (Figure 7B) secreted by WEHI-231 cells. EVs with ectosomal markers (CD9+ANXA2+) were approximately four-times more abundant than EVs with exosomal markers (CD63+LAMP-1+) (Figure 7C-D). The levels of CD63+LAMP-1+ exosomes did not change regardless of inhibitor treatment or CD24 stimulation (Figure 7D). Interestingly, it appeared that CD63 on its own labeled the EV fraction that did not change while LAMP-1 expression was also likely to be found on CD9+ANXA2+ EVs (Supplemental Figure S4). Surprisingly, we found that inhibition of ROCK, with Y27632, or aSMase, with imipramine or ARC39, potentiated the release of total, GFP+, and CD9+ANXA2 ectosomes when CD24 was stimulated (Figure 7A-D). Inhibition of PI3K, with LY294002, potentiated CD24-mediated release of GFP+ EVs (Figure 7B). Overall, the statistical analysis showed an overall effect of CD24 stimulation on ectosome release (P<0.001) but not exosome release (P=0.18), when analyzing the double positive populations.

**Figure 7:**
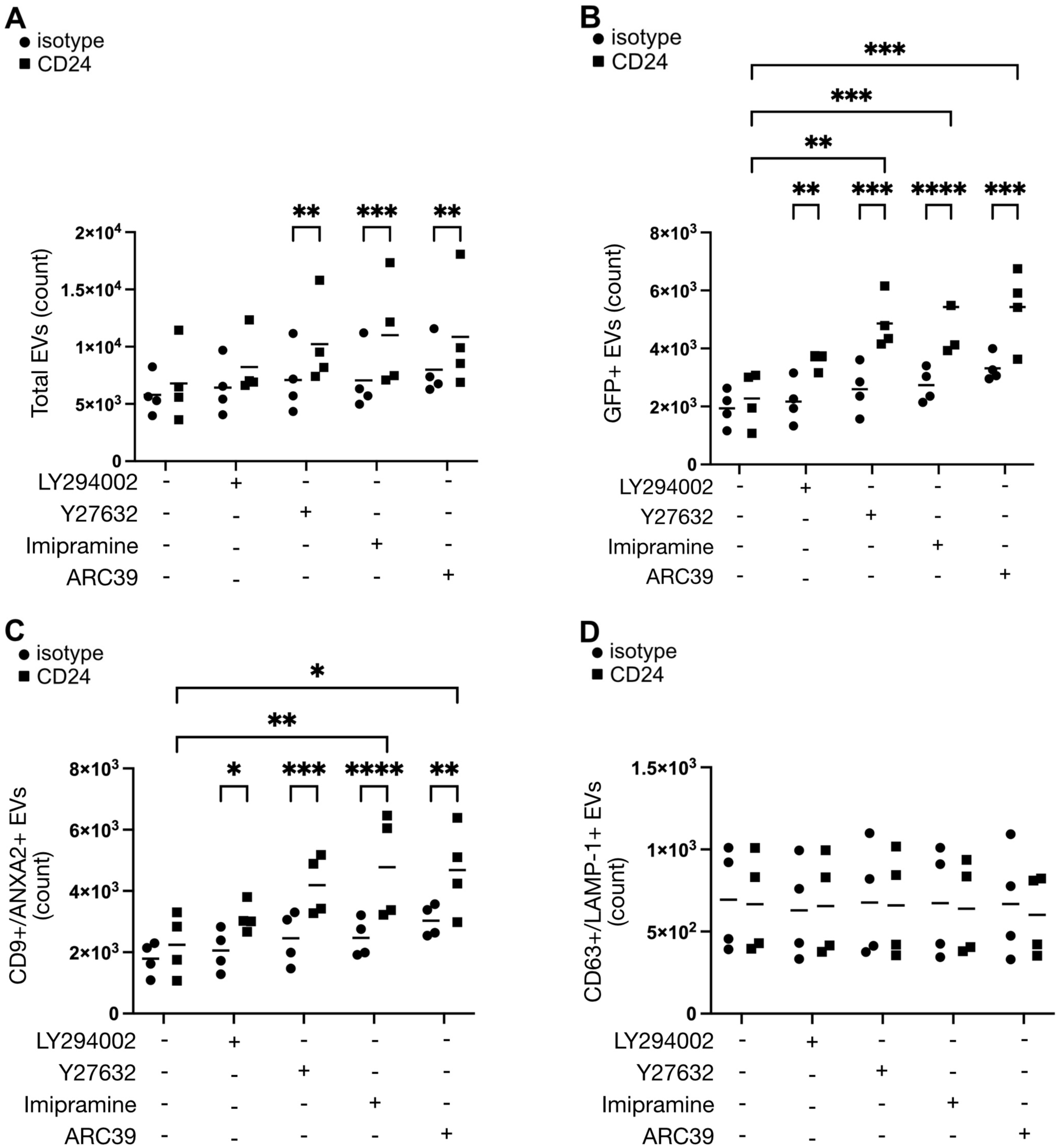
CD24 does not change overall numbers of secreted EVs while PI3K, ROCK and aSMase increase ectosome production when CD24 is stimulated. WEHI-231-GFP cells were pre-treated with LY194002, Y27632, imipramine, or ARC39 for 15 min, then stimulated with anti-CD24 (CD24) or isotype control (isotype) for 15 mins, followed by a washout and 1hr incubation in EV-depleted media. EVs were stained with (A) membrane stain vFRed, (B) vFRed and GFP, (C) vFRed and ectosomal markers (CD9 and ANXA2), or (D) vFRed and exosomal markers (CD63 and LAMP-1). n=4, statistical significance determined by a two-way ANOVA, followed by Sidak’s multiple comparison test *p < 0.05, **p < 0.01, ***p < 0.005, ****p < 0.001. (A) Stimulation significant at P < 0.0001, (B) Stimulation significant at P < 0.0001 and inhibitor treatment significant at P < 0.01, (C) Stimulation significant at P < 0.0001, and (D) Stimulation and inhibitor treatment were not significant.

The size of the EVs need not change with any of the treatments (Supplemental Figure S5). Thus, CD24 stimulation in combination with suppression of ROCK or aSMase significantly increases the secretion of ectosomes but not exosomes. These data suggest that the pathway identified above (CD24/aSMase/PI3K/ROCK/mTORC2/actin) regulates the packaging of cargo that allows uptake of EVs by recipient cells as opposed to regulating total EV release. Moreover, when this packaging is inhibited ectosome secretion increases, which suggests that packaging of cargo normally reduces the rate of total ectosome production.

Lastly, we performed live cell imaging to visualize membrane dynamics in response to CD24 stimulation. We clearly saw that the plasma membrane labeled with GFP displayed a substantial increase in dynamic movement when cells were stimulated through CD24 (Figure 8 and Supplemental Video Files 1-4). In some cases, we were able to visualize EV-like structures appearing just outside the cell or as bubbles associated with the plasma membrane (Figure 8A, inset); however, this was extremely difficult to quantify as the structures were very transient.

**Figure 8.**
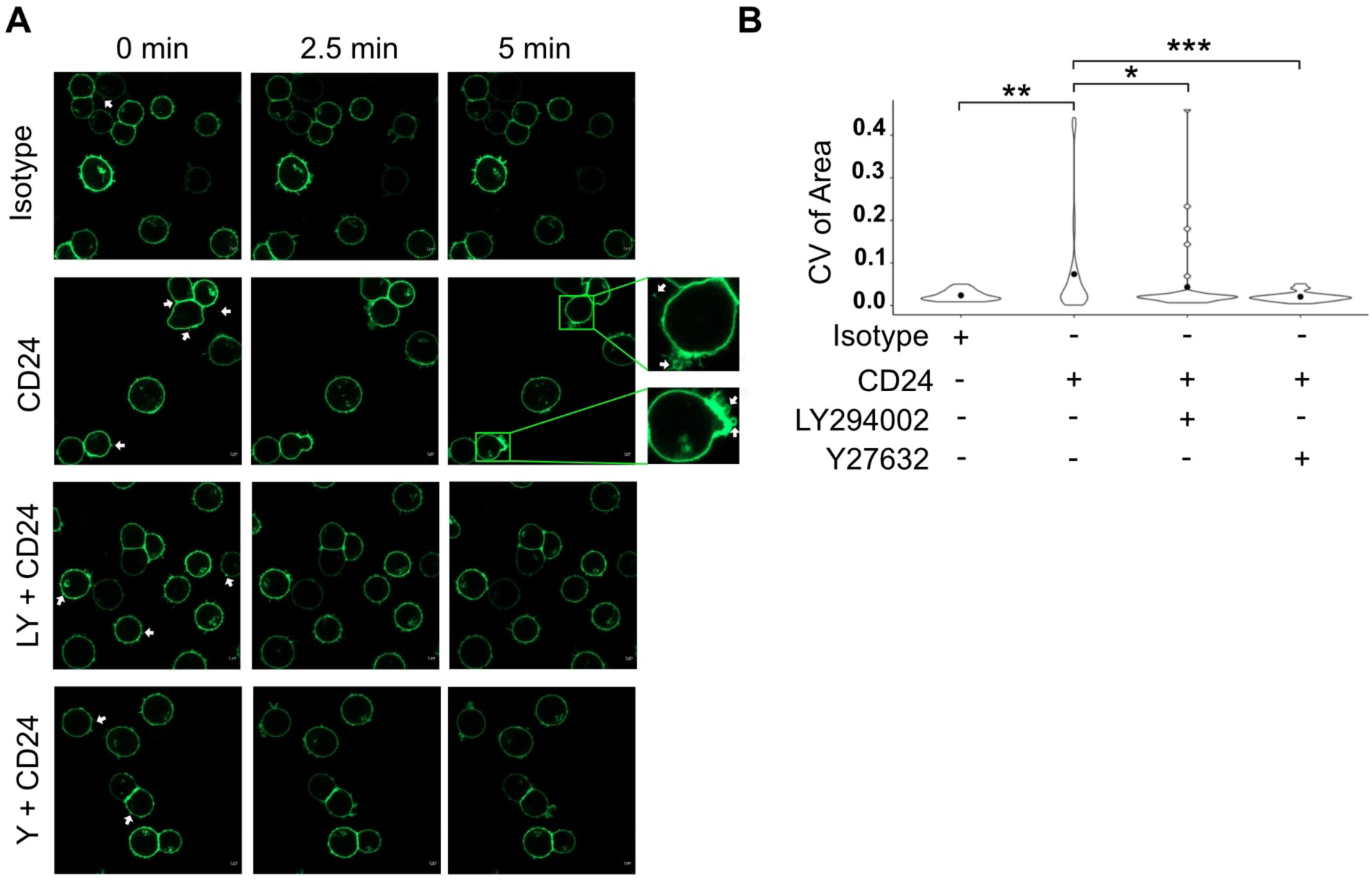
CD24 induces dynamic membrane movement in B cells via PI3K and ROCK. WEHI-231-GFP cells were pre-treated with LY294002 (CD24+LY) or Y27632 (CD24+Y) or DMSO for 15 min, then stimulated with anti-CD24 (CD24) or isotype (Iso) control. (A) The cells were captured at 30 min after stimulation. White arrows indicate where EV release was seen in the videos. Scale bar = 2 μm. (B) Cell activation was determined using the coefficient of variance (cv) of the area over 10 images covering 5 min and visualized with violin plots (the filled black circle indicates the mean). Five to 12 cells were analyzed per video for a total of 38-40 cells analyzed per treatment group. Pairwise comparisons using the Kolmogorov-Smirnov test were used to determine significant differences *P<0.05, **P<0.01.

Using the coefficient of variance of the area of each cell over time, as a measure of changes to cell membrane shape and size, which is an indicator of membrane activation, we found that CD24 significantly increased the dynamic changes to cell shape and size while inhibition of PI3K or ROCK significantly inhibited this increase (Figure 8B), suggesting that membrane dynamics contribute the generation of bioactive ectosomes.

## Discussion

We have identified a novel pathway regulating the formation of bioactive EVs that can be taken up by recipient cells, namely the aSMase/PI3K/mTORC2/ROCK/actin pathway. By exploiting bioinformatics pathway analysis, we were able to identify PI3K and mTOR as potential nodes regulating CD24-mediated bioactive EV formation. Further elucidation of potential regulators allowed us to determine that this pathway is initiated by aSMase which, through an unknown mechanism, activates PI3K to promote actin rearrangement downstream of the mTORC2/ROCK axis. To the best of our knowledge, this is the first time that CD24 has been linked to PI3K and to aSMase. Interestingly, via single EV analysis, we found that similar to previous work^37, 39^, CD24 did not increase total EV numbers. Thus, these data suggest that activation of the aSMase/PI3K/mTORC2/ROCK/actin pathway by CD24 regulates the formation of a subset of EVs that can be taken up by recipient cells but not EV formation *per se*. Lastly, we found that PI3K and ROCK regulate membrane dynamics that correlates with the generation of these bioactive EVs.

Differentiation of exosomes from ectosomes is difficult as there are no markers that distinguish these with 100% certainty^2^. However, we believe that CD24 induces the release of ectosomes that are pinched off the plasma membrane. The supporting evidence comes from previous publications in addition to the data we show here. Firstly, in multiple instances, we found the size of the EVs released by these cells to be predominantly in the 100-180 nm range, which is more in line with the size of ectosomes^37–39^. Next, by TEM we found no evidence of multi-vesicular bodies in the donor cells but clear evidence of EVs in the extracellular environment^37^. We found that surface CD24 protein, as detected by antibody, is exchanged between B cells, suggesting that these EVs are formed from the surface of the cell^37, 39^. In this study, GW4869, an exosome biogenesis inhibitor, had no effect on CD24-mediated EV release whereas impramine, an ectosome inhibitor, significantly inhibited EV release^60^. Moreover, the involvement of actin in CD24-mediated EV release also suggests an ectosomal source rather than via the endocytic pathway that generates exosomes. Lastly, here we find that EVs bearing ectosome markers were more abundant than EVs with exosomal markers released from these cells. There was increased CD24-mediated secretion of ectosomal EVs in the presence of key inhibitors. Overall, these data suggest that CD24 primarily regulates packaging of cargo into ectosomes, not exosomes.

This study revealed a range of pharmacological agents that can effectively inhibit the formation of bioactive EVs that specifically carry GFP and IgM and that are subsequently taken up by recipient cells. Inhibitors of the PI3K/Akt pathway, LY294002 and MK2206, respectively inhibited bioactive EV generation. In addition, used Torin1, and JR-AB2-011 but not rapamycin regulated the transfer of EVs. These data showed that mTORC2, but not mTORC1, is essential for CD24-mediated EV transfer. The ROCK inhibitor Y27632, which can regulate multiple cellular processes such as cell growth, differentiation, and cytoskeleton regulation also inhibited bioactive EV formation ^61^. Cytochalasin D, which disrupts actin filaments of the cytoskeleton, particularly inhibiting actin polymerization blocked EV transfer. Lastly, inhibitors of aSMase (impramine and ARC39) effectively blocked EV transfer.

Contrary to previous reports^13, 31^, we did not find any involvement of intracellular calcium in the regulation of EV release in these cells (Supplemental Figure S3B).

Lipid rafts play a key role in the synthesis and function of EVs^62^. Ceramide-enriched lipid rafts have also been found in exosomal membranes^63^. Herein, we report decreased levels of EV transfer in the presence of imipramine or ARC39, which prevents the activation of aSMase, and the subsequent increase in ceramide levels. Thus, in these cells, aSMase may be inducing the formation of ceramide-enriched lipid rafts, which in turn can promote the membrane curvature needed for EV formation^64^ or regulate loading of specific cargo^65^. Moreover, alterations to membrane thickness can regulate the incorporation of proteins into microdomains^66^, supporting a mechanism by which changes to local ceremide concentrations could alter cargo packaging. The increase in local ceramide concentrations might also displace cholesterol in the lipid rafts to alter their function and initiate signal transduction^67^. However, alterations in lipid composition in lipid rafts in response to CD24 stimulation have yet to be determined.

Previous work has linked CD24 to the recruitment of PTEN, a negative regulator of PI3K signaling, to the plasma membrane to induce autophagy by inhibition of downstream proteins Akt and mTORC1^68^. However, this is opposite to what we found, which was an activation of PI3K by CD24.

CD24 has been found to mediate signal transduction by recruiting Src family protein tyrosine kinases, including Fgr, Lyn, and Lck to lipid rafts^35, 36^. Specifically, Lyn is activated by CD24 in B cells^35^. Lyn activates PI3K in colorectal cancer cells and myeloma cells^69, 70^. In endometrial cells, Lyn has been shown to be activated by long chain glucosylceramide and in neutrophils by lactosylceramide^71, 72^. Therefore, activation of Lyn by re-organization of lipid raft components due to the increase in ceramide, as discussed above, may promote the activation of PI3K downstream of CD24.

This is the first report of regulation of membrane dynamics by CD24, to the best of our knowledge. We observed a clear increase the number of cells with significant changes to cell area over time, which was inhibited by PI3K and ROCK inhibitors. PI3K can regulate membrane attachment to the actin cytoskeleton by decreasing the levels of (PI(4,5)P_2_)^20, 21^, which can allow increased movement of the plasma membrane. ROCK, on the other hand, can directly regulate actin dynamics, which in turn regulates ectosome formation^45^. The relationship between membrane dynamics and packaging of specific cargo to induce formation of bioactive ectosomes is an area for future work.

Interesting, inhibition of ROCK and aSMase potentiated the release of EVs enriched in ectosomal markers when CD24 was stimulated. This was unexpected as EV transfer was clearly inhibited with the same treatments. These data suggest that ROCK and aSMase promote the loading of cargo, presumably a protein, that promotes the uptake of these bioactive EVs by recipient cells. Furthermore, the data suggest that in the absence of this packaging, there is an increase in the formation of ectosomes that are not taken up by recipient cells, as there is an increase in ectosome number but not an increase in GFP^+^IgM^+^ recipient cells. Future work will focus on identifying the changes in surface EV cargo that regulates EV uptake by recipient cells and that is modified by CD24 stimulation. Moreover, future work on the rate of ectosome synthesis in the presence and absence of cargo loading may reveal additional mechanisms regulating ectosome formation.

The function of CD24-mediated EV transfer in the bone marrow is not clear. However, our previous data showed that these EVs transfer functional receptors, including CD24 and the B cell receptor (BCR) to recipient B cells^39^. The transfer of these receptors increased apoptosis in the recipient cells. Therefore, it is possible that increased activation of the CD24 signaling pathway would cause transfer of the pro-apoptotic function of CD24 in recipient cells causing apoptosis in bystander cells. We hypothesize that this may be a mechanism to regulate homeostasis whereby increased numbers of CD24-positive cells, which increases in the C/C’ stage of B cell development^39^, would result in release of more of these bioactive EVs, which, in turn, would reduce the number of bystander B cells and maintain B cells numbers at a steady-state.

Identification of the trigger for regulating homeostatic proliferation in this manner is a current area of investigation as the ligand for CD24 on B cells remains unknown. Nevertheless, the data presented here suggest that interruption of the aSMase/PI3K/mTORC2/ROCK/actin could block the transfer of functional pro-apoptotic receptors, which may increase the numbers of developing B cells leading to autoimmune or lymphoproliferative disorders; a hypothesis that we will be addressing in the future.

Interestingly, the PI3K pathway is essential for proliferation and development of bone marrow B cells downstream from the pre-BCR^73^, which is in contrast to the pro-apoptotic function of CD24 in these cells^35, 37^. Therefore, in the case of CD24, it is likely that the balance of pro-apoptotic pathways (ie. p38) outweighs the activation of pro-proliferative pathways (ie. ERK and PI3K)^74^ whereas the opposite balance occurs with pre-BCR signaling. We previously observed that engagement of the BCR in WEHI-231 cells, composed of IgM and Igα/β, also induces EV-mediated transfer of functional proteins, suggesting that multiple receptors could use the PI3K pathway to regulate EV transfer, thus, revealing a new role for PI3K in cellular function. Future investigations should determine if the PI3K pathway is a common regulator for bioactive EV production from B cells.

In summary, here we provide compelling evidence that CD24 activates an aSMase/PI3K/mTORC2/ROCK/actin signaling pathway to induce the formation of bioactive EVs that carry functional cargo to recipient cells. Importantly, these findings provide a new avenue for research into the regulation of ectosome generation.

## Acknowledgements

The authors wish to thank the Medical Laboratories at Memorial University for their support with flow cytometry and confocal imaging.

**Supplemental Files 1-4. Videos of cell activation in response to CD24 after 30 min in the presence or absence of LY294002 or Y27632.**

