## Supplementary material for "CD24 regulates the formation of ectosomes in B lymphocytes": Supplemental data.docx

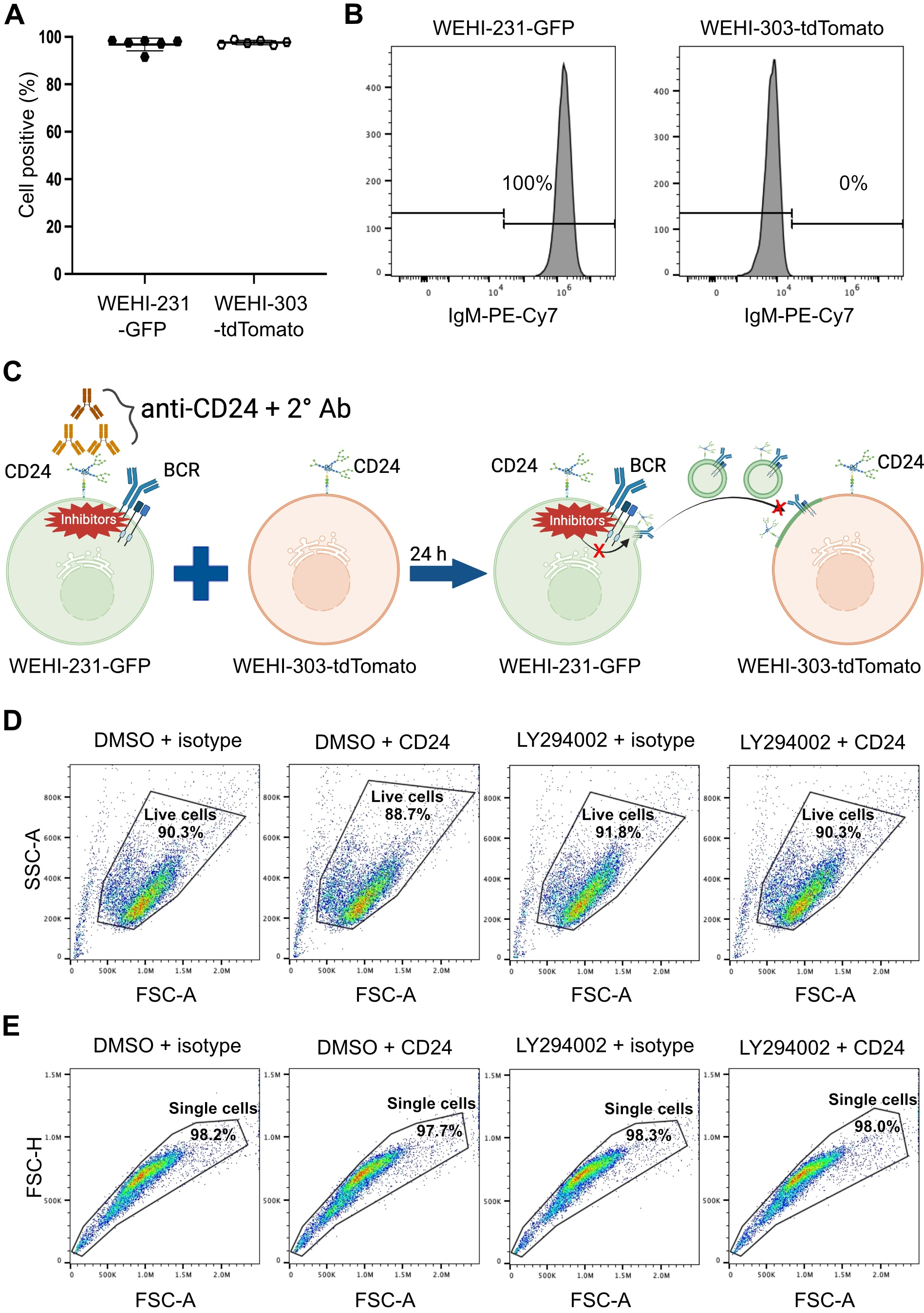

**Supplemental Figure S1. Pre-treatment with inhibitor on donor WEHI-231-GFP cells inhibits the transfer of GFP and IgM from donor to recipient cells in response to CD24.** (A) The percentage of cells positive for GFP and tdTomato. (B) WEHI-231-GFP are 100% IgM positive while WEHI-303-tdTomato cells do not express IgM on their cell membrane. (C) Schematic of experimental design of model system. WEHI-231-GFP cells were pre-treated with a chemical, for example, LY294002 or DMSO, for 15 min, then stimulated with isotype control or anti-CD24 for 15 min, followed by washout and then a 24 h co-culture with WEHI-303-tdTomato cells. GFP and IgM are then detected with flow cytometry. (D) Representative dotplots of live cell population sorting based on SSC-A/FSC-A. (E) Representative dotplots of the single cell population sorting based on FSC-H/FSC-A.

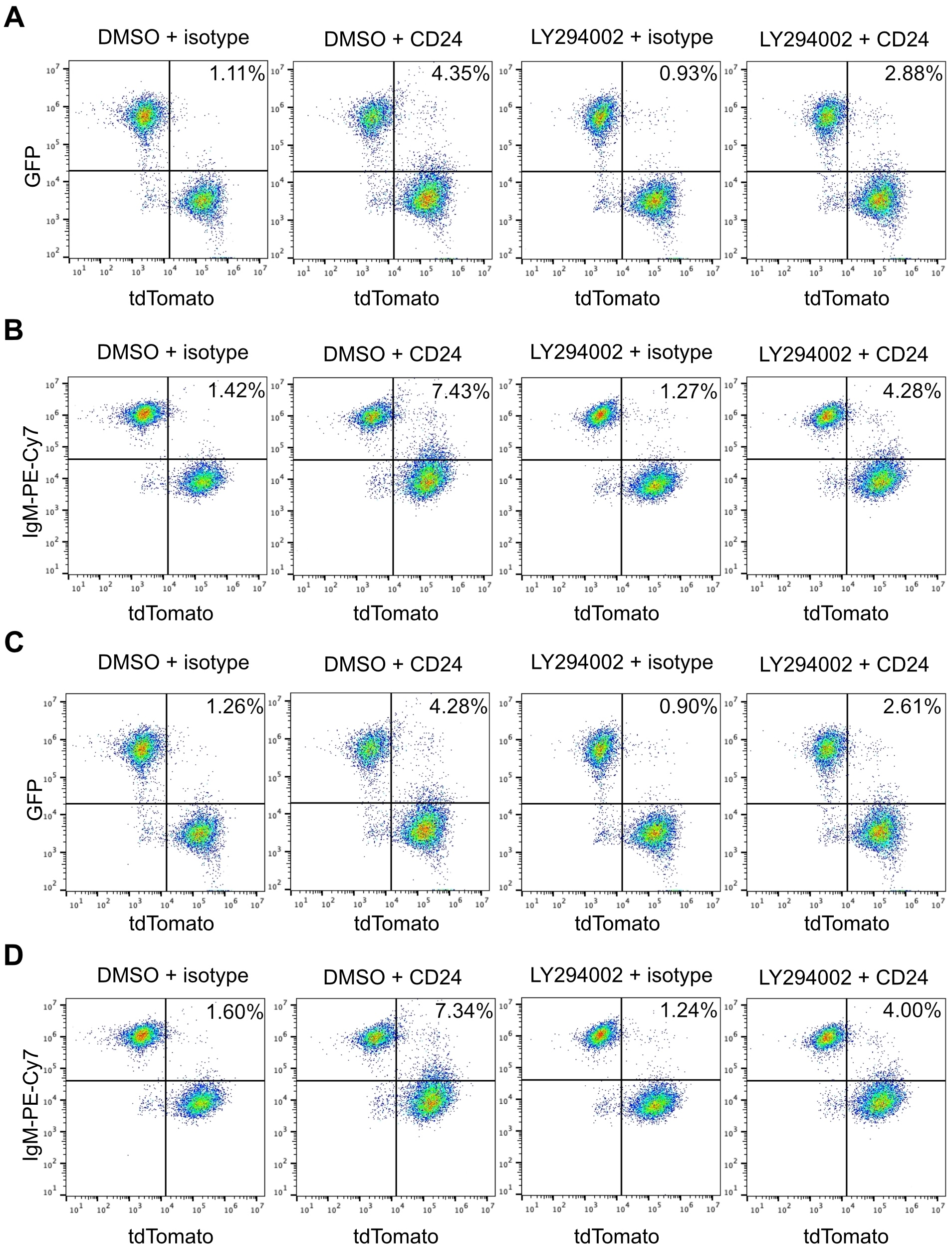

**Supplemental Figure S2. Live cell and single cell gating analyses show the same results for transfer of GFP and IgM.** Representative dotplots of GFP-positive (A) and IgM-positive (B) on WEHI-303-tdTomato cells after co-culture with stimulated donor WEHI-231-GFP cells based on live cell population sorting as shown in Figure S1D. Representative dotplots of GFP-positive (C) and IgM-positive (D) on WEHI-303-tdTomato cells after co-culture with stimulated donor WEHI-231-GFP cells based on single cell sorting as shown in Figure S1E.

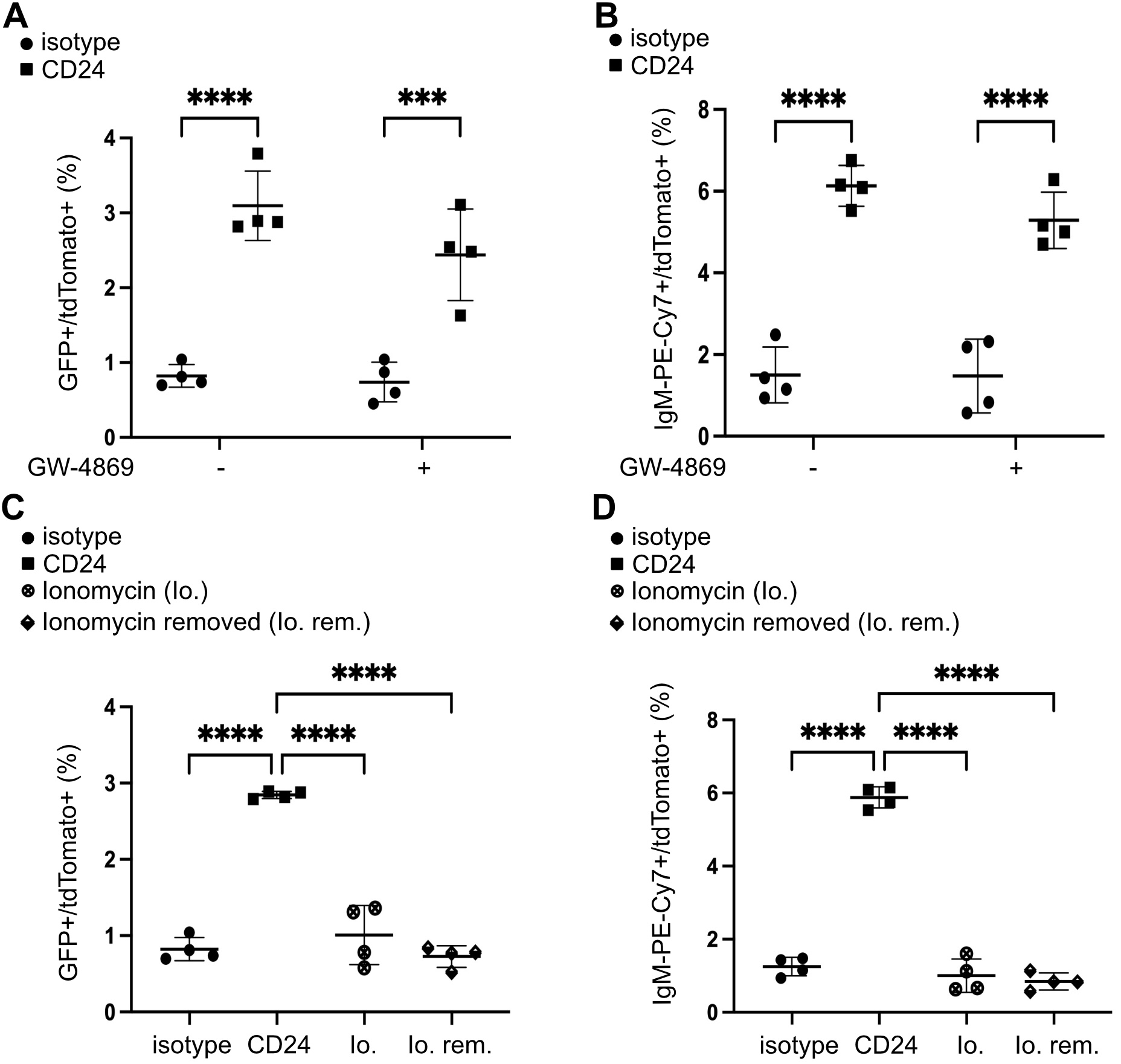

**Supplemental Figure S3. Neither nSMase nor calcium regulates EV release.** (A-B) WEHI-231-GFP cells were pre-treated with GW4869 or DMSO for 15 min, then stimulated with anti-CD24 or isotype control for 15 min, followed by washout and then a 24 h co-culture with WEHI-303-tdTomato cells. (A) Percent GFP and tdTomato double-positive cells and (B) Percent IgM and tdTomato double-positive cells after 24 h incubation. n=4, statistical significance determined by a two-way ANOVA (interaction significant at P=0.2669 for A and P=0.4195 for B) followed by the Sidak’s multiple comparison test ***P<0.005, ****P<0.001. (C-D) WEHI-231-GFP cells were stimulated with isotype control, anti-CD24, ionomycin or ionomycin removed after 15 min, followed by 24 h co-culture with WEHI-303-tdTomato cells. (C) Percent GFP and tdTomato double-positive cells. (D) Percent IgM and tdTomato double-positive cells after 24 h incubation. n=4, statistical significance was determined by a one-way ANOVA followed by the Sidak’s multiple comparison test ****P<0.001.

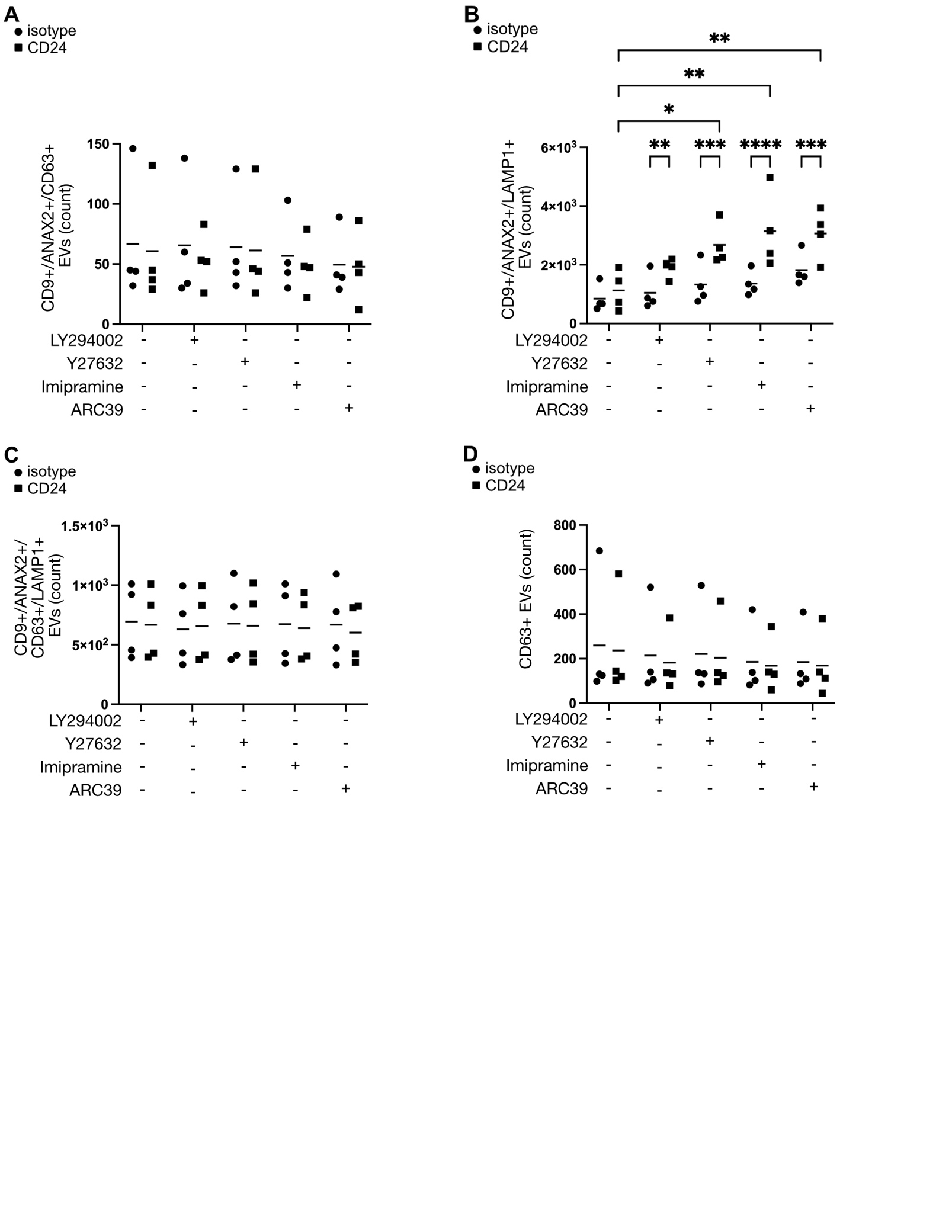

**Supplemental Figure S4:** **Exosomal EV detection is driven by CD63 but not LAMP1.**

WEHI-231-GFP cells were pre-treated with LY294002, Y27632, imipramine, or ARC39 for 15 min, then stimulated with anti-CD24 (CD24 EVs) or isotype control (isotype EVs) for 15 mins, followed by a washout and 1hr incubation in EV-depleted media. EVs were isolated by centrifugation, then stained with membrane stain vFRed, and the markers indicated. Data are represented as individual points of mean, n=4, statistical significance determined by a two-way ANOVA, followed by Sidak’s multiple comparison test. Statistical significance is indicated as follows: *p < 0.05, **p < 0.01, ***p < 0.005, ****p < 0.001. (A) No significant differences in exosome populations were observed, (B) There was a significant effect of stimulation (P < 0.0001) and inhibitor treatment (P < 0.01), (C) No significant effect of stimulation or inhibitor treatments. (D) No significant effect of stimulation or inhibitor treatments.

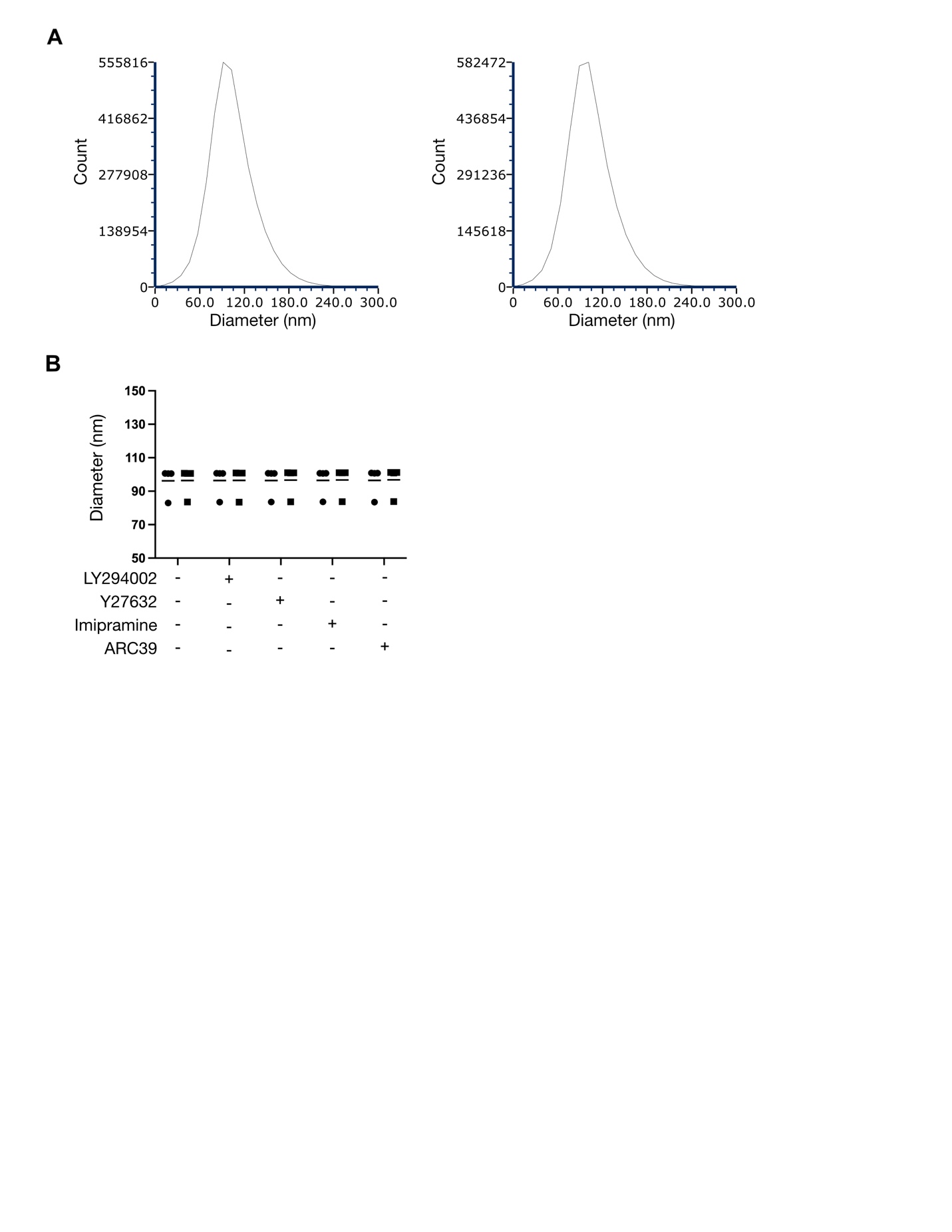

**Supplemental Figure S5: The average diameter of total EVs does not change with CD24 stimulation or treatment with inhibitors.** WEHI-231-GFP cells were pre-treated with LY294002, Y27632, imipramine, or ARC39 for 15 min, then stimulated with anti-CD24 (CD24 EVs) or isotype control (isotype EVs) for 15 mins, followed by a washout and 1hr incubation in EV-depleted media. EVs were isolated by centrifugation, then stained with membrane stain vFRed. (A) Representative histograms of vFRed positive EV diameters from isotype control (left panel), and CD24 stimulated (right panel) conditions. (B) Data are represented as individual points of mean, n=4, statistical significance determined by a two-way ANOVA, followed by Sidak’s multiple comparison test. Stimulation and inhibitor treatment were not significant.
